# Predictable fluctuations in excitatory synaptic strength due to natural variation in presynaptic firing rate

**DOI:** 10.1101/2022.04.14.488362

**Authors:** Naixin Ren, Ganchao Wei, Abed Ghanbari, Ian H. Stevenson

## Abstract

Many controlled, *in vitro* studies have demonstrated how postsynaptic responses to presynaptic spikes are not constant but depend on short-term synaptic plasticity (STP) and the detailed timing of presynaptic spikes. However, the effects of short-term plasticity (depression and facilitation) are not limited to short, sub-second timescales. The effects of STP appear on long timescales as changes in presynaptic firing rates lead to changes in steady-state synaptic transmission. Here we examine the relationship between natural variations in the presynaptic firing rates and spike transmission *in vivo*. Using large-scale spike recordings in awake mice from the Allen Institute Neuropixels dataset, we first detect putative excitatory synaptic connections based on cross-correlations between the spike trains of millions of pairs of neurons. For the subset of pairs where a transient, excitatory effect was detected, we use a model-based approach to track fluctuations in synaptic efficacy and find that efficacy varies substantially on slow (∼1 minute) timescales over the course of these recordings. For many connections, the efficacy fluctuations are correlated with fluctuations in the presynaptic firing rate. To understand the potential mechanisms underlying this relationship, we then model the detailed probability of postsynaptic spiking on a millisecond timescale, including both slow changes in postsynaptic excitability and monosynaptic inputs with short-term plasticity. The detailed model reproduces the slow efficacy fluctuations observed with many putative excitatory connections, suggesting that these fluctuations can be both directly predicted based on the time-varying presynaptic firing rate and, at least partly, explained by the cumulative effects of STP.

## Introduction

The firing activity of individual neurons can vary over time due to external stimuli (Vogels et al., 2005), the animal’s behavior (Churchland et al., 2012; Shenoy et al., 2013), and the internal state of the network (Poulet and Petersen, 2008; McGinley et al., 2015). Although many statistical models assume that coupling between neurons has a fixed strength (Harris et al., 2003; Truccolo et al., 2005; Pillow et al., 2008), firing rate variation can directly affect synaptic strength due to short-term synaptic plasticity (STP). For instance, synapses with short-term depression should be weaker when presynaptic firing rates are higher and stronger when the rates are lower. On the other hand, synapses with short-term facilitation should be stronger when presynaptic firing rates are higher and weaker when the rates are lower. In many studies, STP is induced *in vitro* by modifying the presynaptic firing rate using external electrical stimulation at different frequencies (Abbott et al., 1997; Tsodyks and Markram, 1997; Fortune and Rose, 2001; Chung et al., 2002; Zucker and Regehr, 2002). These studies illustrate how STP can act to high-pass or low-pass filter a neuron’s inputs, and could potentially underlie working memory and decision-making processes (Deng and Klyachko, 2011).

One prediction from these *in vitro* studies of STP using simple, controlled stimuli is that synaptic strengths should also vary in response to the natural changes in firing rate that occur *in vivo*. Indeed, slice studies have shown that, for neurons receiving naturalistic spike trains as input, synaptic strength varies substantially over time, with patterns matching those predicted by STP (Klyachko and Stevens, 2006; Kandaswamy et al., 2010; Costa et al., 2013). Studies in awake animals have also shown that postsynaptic responses to presynaptic spikes vary depending on the preceding presynaptic inter-spike intervals (Swadlow and Gusev, 2001; Stoelzel et al., 2008, 2009; English et al., 2017) and presynaptic firing rates (Fujisawa et al., 2008; Stoelzel et al., 2015; McKenzie et al., 2021). Together, these findings suggest a general conclusion that the natural variation in presynaptic firing rates *in vivo* may be accompanied by large fluctuations in synaptic strength and that these changes may be due to the effects of STP on longer timescales.

Quantifying synaptic strength *in vivo* is a difficult statistical problem. Many studies of synaptic transmission have used intracellular recordings to study the dynamics in individual postsynaptic potentials or currents (Pala and Petersen, 2015, 2018; Sedigh-Sarvestani et al., 2017). Long-term extracellular spike recordings may allow us to track slow fluctuations more easily *in vivo*. Unlike intracellular signals that contain postsynaptic responses to single presynaptic spikes, extracellular spikes are sparse binary events, and studies of synapses often rely on the cross-correlograms (CCGs) between the pre- and postsynaptic spike trains. If two neurons are connected with an excitatory synapse, there is often a fast-onset, short-latency peak in the CCG (Perkel et al., 1967; Fetz et al., 1991), and several techniques have been developed to automatically detect these connected pairs (Barthó et al., 2004; Amarasingham et al., 2012; Kobayashi et al., 2019; Ren et al., 2020).

Synaptic transmission with spikes is often measured by “synaptic efficacy” using the CCG, which is defined as the excess probability of postsynaptic spiking in the short transmission window following presynaptic spikes (Levick et al., 1972). By focusing on specific presynaptic spike patterns, such as pairs with specific inter-spike intervals (ISI), previous studies have shown how synaptic efficacy can vary depending on features of presynaptic spike timing (Usrey et al., 2000; Swadlow and Gusev, 2001; English et al., 2017). Modeling individual spike transmission probabilities for the entire spike train can also reveal how presynaptic spike features, as well as different types of plasticity, interact (Ghanbari et al., 2017, 2020; Song et al., 2018; Wei and Stevenson, 2021). Together, descriptive and model-based approaches can track synaptic dynamics *in vivo* (Swadlow and Gusev, 2001; English et al., 2017; Ghanbari et al., 2017, 2020) and reveal changes associated with learning, behavior, or stimuli (Fujisawa et al., 2008; Ghanbari et al., 2020; McKenzie et al., 2021). Here we focus on a general description of the ongoing changes in synaptic strength that may be due, merely, to changes in presynaptic rate. We divide long-term spike recordings into windows to measure the fluctuations in synaptic efficacy on a timescale of minutes and model the changes as a function of presynaptic ISI to build the potential link between these slow changes and STP.

Using *in vivo* large-scale spike recordings from the Allen Institute Neuropixels dataset (Siegle et al., 2021), we examine natural fluctuations in synaptic efficacy and firing rates and quantify the relationships between them. These recordings, from awake, behaving mice, contain the spiking of hundreds of neurons from many brain regions while the mice view natural and artificial visual stimuli (Siegle et al., 2021). Since neurons in different brain regions have different firing rates and firing regularity (Mochizuki et al., 2016), we also examine whether putative synapses from different regions show different patterns of efficacy fluctuations. We find that, for individual connections, the synaptic efficacy often varies substantially over the course of the recording. Interestingly, the efficacy is often correlated with the presynaptic firing rate, with the postsynaptic firing rate, or both. We then test if the time-varying efficacy is predictable based on the presynaptic firing. We model the time-varying synaptic effect as a function of presynaptic inter-spike interval (ISI) by fitting a Generalized Bilinear Model (GBLM) to the postsynaptic spike trains. For many connections, the GBLM reproduces the observed efficacy fluctuations, suggesting that these fluctuations can be, at least partly, explained by a cumulative effect of STP. Altogether, these results illustrate how natural fluctuations in firing rate could lead to large, but predictable, changes in synaptic strength *in vivo* and in many different brain areas.

## Methods

Here we use the Visual Coding – Neuropixels dataset from the Allen Institute (Dataset: Allen Institute MindScope Program (2019). Allen Brain Observatory -- Neuropixels Visual Coding [dataset]. Available from brain-map.org/explore/circuits). During the recordings, head-fixed mice viewed a standardized set of visual stimuli (including Gabor patches, full-field drifting gratings, moving dots, and natural images and movies) while they were free to run on a wheel. We analyze 20 recordings here (ecephys_session_id: 715093703 – 746083955, 819186360 – 847657808), 6 from female mice (125 ± 11.6 days), 14 from male mice (117.8 ± 8.3 days). 9 recordings were from wild-type mice, while the remainder were transgenic Cre lines expressing channelrhodopsin in inhibitory interneuron subtypes (Sst-IRES-Cre; Ai32, Pvalb-IRES-Cre; Ai32 and Vip-IRES-Cre; Ai32). Each recording lasts approximately 2.5 hours and contains 365-893 well-isolated single units from 5 or 6 Neuropixels probes from mouse visual cortex, thalamus, and hippocampus. In 10 recordings, the mice had a resting period that last for about 30 minutes. This period is identified as spontaneous activity period for further analysis. Additional information about this dataset can be found in Siegle et al. (2021).

### Detection of putative excitatory synaptic connections

We detect putative synaptic connections using the cross-correlograms (CCG) between pairs of neurons: a source neuron (presynaptic) *i* and target neuron (postsynaptic) *j*. For the binned spike trains of the two neurons, *n*_*i*_ and *n*_*j*_ (1 when there is a spike and 0 otherwise), the CCG is given by *y*_*ij*_(*m*) = ∑_*t*_ *n*_*i*_(*t*)*n*_*j*_(*t* − *m*), where *m* denotes the interval between pre- and postsynaptic spikes, and *y*_*ij*_(*m*) is the number of the times spikes in *n*_*i*_ and *n*_*j*_ are separated by an interval 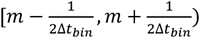 for binsize Δ*t*_*bin*_ (1 ms used here).

Given the CCG, we detect putative excitatory synapses in two stages. First, we use a hypothesis test based on the jittered spikes to detect fast changes in the CCG, and then, we use a model-based approach to estimate and screen for putative synaptic effects at specific latencies and timescales. Our initial screening of neuron pairs is based on comparing the observed CCG to a null model of the CCG generated using jittered spike times. An exact null distribution can be approximated by directly sampling jittered spike trains. However, here, for computational convenience we approximate the null distribution using the simplifying assumption that the bins of the CCG are independent (Amarasingham et al., 2012). We assume that *y*_*ij*_(*m*)*∼Binomial*(*n*_*pre*_, *µ*(*m*)/*n*_*pre*_) where *n*_*pre*_ is the total number of presynaptic spikes and *µ*(*m*) = (*y*_*ij*_ ∗ *f*)(*m*), the convolution of the CCG with a jitter function. Here we use a 10ms uniform jitter *f*(*t*) = 1 for |*t*| < 5*ms*, 0 otherwise. We then compute p-values for each bin of the CCG using the cumulative distribution function (CDF) of the binomial distribution and evaluate statistical significance correcting for multiple comparisons by controlling the false discovery rate using a Benjamini–Hochberg procedure (*α* = 0.00001). We then identify pairs of neurons where two criteria are satisfied: 1) there are two neighboring statistically significant bins in the CCG (where m>0), and 2) the center bin (*m* = 0) is not statistically significant. This test yields 0.5% of pairs of neurons that have fast, transient changes in their CCGs.

For each pair of neurons that satisfies the testing criteria we then fit a parametric model that aims to estimate the latency, timescale, and strength of a putative synaptic effect. Here we fit an extended GLM based on the model used in Ren et al., (2020). Briefly, we describe the CCG using two components: 1) a slow fluctuation due to background fluctuations and 2) a fast, transient effect due to the synaptic effect. The rate of spike counts *λ* at bin *m* in a CCG is given by:

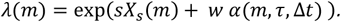

Here *sX*_*s*_ describes the slow fluctuation, where *X*_*s*_ represents a set of smooth basis functions learned by applying a low-rank, nonlinear matrix factorization to all the cross-correlograms in the dataset (Ren et al., 2020). And *wα*(*m, τ*, Δ*t*) represents the fast synaptic effect, where *w* is the synaptic strength and *α*(*m, τ*, Δ*t*) is an alpha function with a time constant *τ* and a latency Δ*t* that is then convolved with the auto-correlogram of the presynaptic neuron to account for possible bursting in the neuron. After fitting the parameters {*s, w, τ*, Δ*t*} using a penalized maximum likelihood method, we select the neuron pairs where the CCG has a strong sharp peak (*w* > 0.3, *τ* < 0.8 *ms*) with a short latency (Δ*t* < 10 ms) and a relatively flat slow fluctuation (the coefficient of variation of the slow fluctuation < 0.15). In the end, we detect 1382 putative excitatory connections (∼ 0.17% of all pairs of neurons). This screening procedure returns a set of neuron pairs whose CCGs are consistent with the biophysics of excitatory synaptic transmission, and which we then characterize in more detail.

### Estimating time-varying synaptic efficacy

Synaptic efficacy reflects the (excess) probability of observing a postsynaptic spike following a presynaptic spike. Here we estimate synaptic efficacy using the model-based approach described in the previous section. Namely, we calculate the efficacy by comparing the model with and without the synaptic effect,

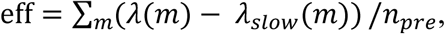

where *λ*_*slow*_ is *λ* evaluated with *θ* = {*s, w* = 0} and *n*_*pre*_ is the number of presynaptic spikes. To estimate changes in observed efficacy over time, we divide the pre- and postsynaptic spike trains into 5-minute windows with 80% overlap and estimate efficacy for each window. We fit the cross-correlogram for each window using the extended GLM with a fixed alpha function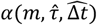, where 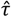 and 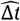 are the estimated parameters from the overall cross-correlogram of the connection.

To examine the relationship between the time-varying efficacy and presynaptic firing rate, we calculate the Spearman correlation coefficient. For comparison, we also shuffle the postsynaptic spike timings to create surrogate data and then calculate the correlations that would be expected by chance. For each presynaptic spike, we resample the subsequent postsynaptic spikes based on the overall CCG. With this shuffling, the overall, average efficacy is preserved, but any fluctuations in the efficacy over long time periods should be due to chance. For each putative synapse, we did the shuffling 100 times and estimate the mean and standard deviation from the shuffled correlations. We then calculate a z-score for the observed correlation to measure how far away it is from the surrogate distribution.

### Modeling short-term synaptic plasticity with spike train observations

In addition to quantifying the observed efficacy from noisy correlograms, we also aim to assess to what extent changes in efficacy can be explained by short-term synaptic plasticity (STP). Here we model the time-varying synaptic effect as a function of presynaptic ISI by fitting a generalized bilinear model (GBLM) to the postsynaptic spike trains. This model has been previously described (Ghanbari et al., 2017; Wei and Stevenson, 2021) and extends previous models of coupled GLMs (Harris et al., 2003; Truccolo et al., 2005; Pillow et al., 2008; Rebesco et al., 2010) to account for fluctuating excitability and plasticity. Briefly, for each connection, we model the postsynaptic firing rate *γ* at time *t* as

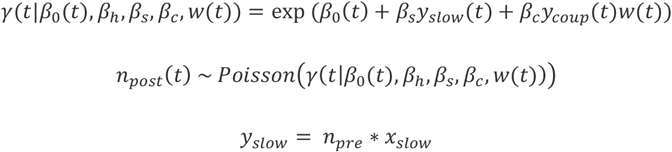

*y*_*coup*_ = *n*_*pre*_ ∗ *x*_*coup*_where *n*_*pre*_ and *n*_*post*_ are the pre- and postsynaptic spike trains, respectively.

Here *β*_0_(*t*) represents a time-varying baseline firing rate, which is estimated using an adaptive filtering algorithm (Wei and Stevenson, 2021). *β*_*s*_*y*_*slow*_(*t*) accounts for slow fluctuations that are shared by the presynaptic neuron and postsynaptic neuron, where *y*_*slow*_ denotes a set of covariates generated by filtering the presynaptic spike train (cubic B spline functions with 4 equally spaced knots over 150 ms *x*_*slow*_). *β*_*c*_*y*_*coup*_*w*(*t*) represents the contribution from the synaptic effect. Here *β*_*c*_*y*_*coup*_(*t*) describes the stationary coupling effect, where *y*_*coup*_ is the presynaptic spike train convolved with the alpha function we learned from the extended GLM *α*(*m, τ*, Δ*t*); and *w*(*t*) describes the effects of STP. Namely, we model *w*(*t*) as a function of presynaptic ISIs Δ*t*_*k*_,

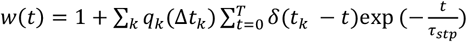

where *q*_*k*_(Δ*t*_*k*_) is a modification function that determines how the synaptic strength increases/decreasing following each presynaptic spike at time *t*_*k*_. We model the modifications using a group of raised cosine basis functions *B*(Δ*t*_*k*_), *q*_*k*_(Δ*t*_*k*_) = *θB*(Δ*t*_*k*_), and assume that the effects of STP decay with *τ*_*stp*_ = 200 ms. We fit the free parameters {*β*_0_(*t*), *β*_*s*_, *β*_*c*_, *θ*} by maximizing the Poisson log-likelihood

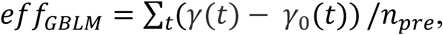

To estimate the parameters, we use an alternating optimization (coordinate ascent): we fit *β*_0_(*t*) using fixed {*β*_*s*_, *β*_*c*_, *θ*} and then fit {*β*_*s*_, *β*_*c*_, *θ*} using fixed *β*_0_(*t*). We repeat this alternating pattern until convergence. After optimization we then have a model-based estimate of the impact of each individual presynaptic spike and can predict how the efficacy would change over time under a model of short-term plasticity. We calculate GBLM estimated synaptic efficacy in a similar way as we estimate efficacy using CCGs,

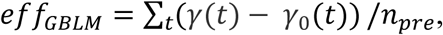

where *γ*_0_ is the postsynaptic rate with the synaptic effect removed *β*_*c*_ = 0. *eff*_*GBLM*_ then reflects the excess probability of observing a postsynaptic spike due to the occurrence of a presynaptic spike within the model.

For comparison, we also fit a static-synapse model where the synaptic coupling effect is *β*_*c*_*y*_*coup*_(*t*) instead of *β*_*c*_*y*_*coup*_(*t*)*w*(*t*). Under the static-synapse model the coupling effect stays the same over time, and will allow us to assess whether STP, specifically, meaningfully improves the description of spike transmission. To quantify how well a model predicts whether there is a postsynaptic spike after each presynaptic spike, we use receiver operating characteristic (ROC) curves, and the area under the curve (AUC) reflects the performance. For each presynaptic spike, we identify a transmission interval (*t*_*a*_, *t*_*b*_) where the coupling filter (alpha function in GBLM) is greater than 0.5, and then we use the model fit within the transmission interval 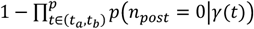 as the score for predicting each individual postsynaptic spike.

## Results

Here we characterize fluctuations in putative synapses from 20 recordings from the Allen Institute Neuropixels dataset (Siegle et al., 2021). We detect putative excitatory synaptic connections based on the CCGs of pairs of neurons using a two-stage synapse detection method – in stage 1, we use a hypothesis test based on the jittered spikes to detect fast changes in the CCG, and, in stage 2, we use a model-based approach to estimate and screen for putative synaptic effects at specific latencies and timescales (see Methods). Figure 1A shows the CCGs of example connections. By using this 2-stage screening procedure, our goal is not to detect as many as putative synapses as possible, but rather to generate a subset of strong putative synapses with CCGs that are consistent with the biophysics of excitatory synaptic transmission, including synaptic latencies (1.29 ± 1.23 s.d. ms, Figure 1B) and time constants (0.55 ± 0.13 s.d. ms, Figure 1C).

**Figure 1.**
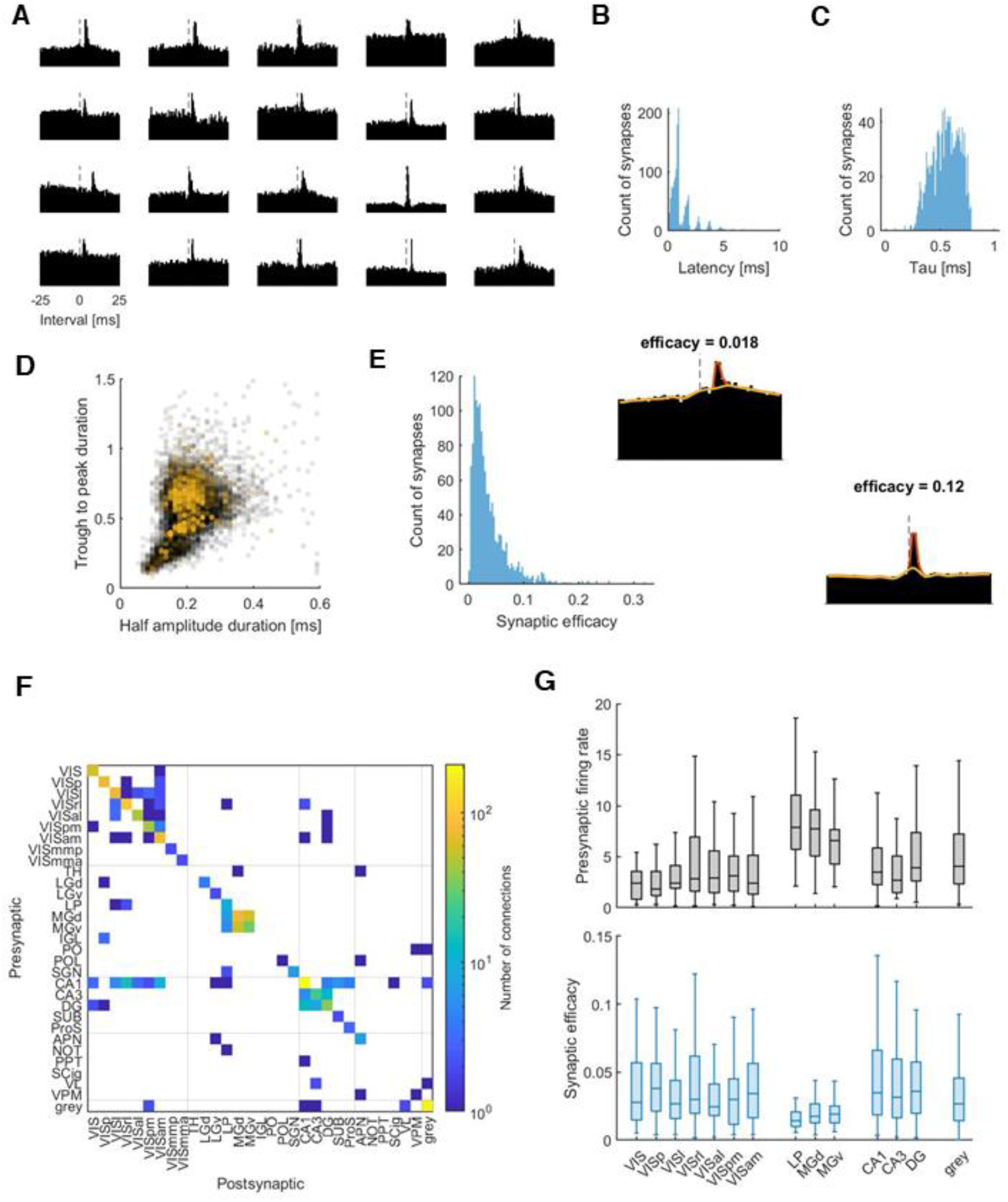
Detection of putative excitatory synaptic connections. A, examples of CCGs of putative excitatory synaptic connections. The bin size is 0.5 ms here for better visualization. In the analysis, we use 1ms. B, the histogram of synaptic latency of all the detected synapses. The distribution tends to peak at 1, 2, 3 … [ms] because the bin size of the CCG is 1 ms. C, the histogram of the synaptic time constant of all the detected synapses. D, two measurements of the duration of waveforms. Each dot represents one neuron. Most putative presynaptic neurons (yellow dots) show broader spike waveforms compared to all the neurons in the dataset (gray dots). E. the histogram of the synaptic efficacy of al the detected synapses. Inserted: an example CCGs with low or high efficacy and the model fits of the extended GLM (red line: model fits with synaptic effect, yellow line: model fits without synaptic effect). F, connectivity matrix across different areas. Y-axis: presynaptic area. X-axis: postsynaptic area. Color represents the number of connections detected (log-transformed). The gridlines separate the brain regions: visual cortex, thalamus, hippocampus, midbrain, and unclassified. G, box plots for the mean firing rate (top) and efficacy (bottom) for brain areas. The areas are grouped by brain regions mentioned above. Only areas with >=10 efferent synapses are shown here. The line inside each box is the median. The top and bottom edges of the box are the upper and lower quartiles, respectively. The whiskers show the nonoutlier maximum and minimum. The outliers are not shown here.

The presynaptic neurons for the majority of putative synapses that we detect have broad waveforms (Figure 1D), which previous work has found tend to correspond to excitatory neurons (Connors and Gutnick, 1990; Barthó et al., 2004; Trainito et al., 2019). For each putative synapse, we estimate the overall synaptic efficacy using an extended GLM (see Methods, Figure 1E). The distribution of synaptic efficacy is approximately log-normal (0.037 ± 0.035, mean ± s.d., Figure 1E), consistent in shape with the previous findings on postsynaptic potentials (Buzsáki and Mizuseki, 2014) and efficacy (English et al., 2017). Here we detect 1383 putative excitatory synapses in total, most of which were between neurons recorded on the same probe (1174 out of 1383) and within the same brain area (1090 out of 1383, Figure 1F).

Across the 20 recordings, we identify putative synapses from many labeled brain regions. The regions with more than 10 efferent synapses included 6 visual cortical areas (primary visual cortex - VISp, lateromedial area - VISl, rostrolateral area - VISrl, anterolateral area - VISal, posteromedial area - VISpm, anteromedial area - VISam), 3 thalamic areas (lateral posterior nucleus - LP, dorsal medial geniculate nucleus - MGd, ventral medial geniculate nucleus - MGv), 3 hippocampal areas (CA1, CA3, and dentate gyrus - DG), and in locations that were not precisely identified (grey or VIS). Synapses from different brain regions have distinct presynaptic firing rates (5.75 ± 9.93 s.d. Hz overall average) and synaptic efficacies (Figure 1G). Notably, putative synapses from thalamic areas appear to have higher presynaptic rates and lower efficacy, on average. Firing rates are expected to fluctuate due to varying stimuli, behavior, and internal state variables. Here we aim to measure the extent to which the efficacy of putative synapses also fluctuates and to determine whether these fluctuations are predictable.

### Presynaptic firing rate and efficacy both fluctuate substantially over time

To examine the natural fluctuations in presynaptic firing rate and synaptic efficacy, we divide the pre- and postsynaptic spike trains into 5-minute windows with 80% overlap and estimate the presynaptic firing rate and efficacy for each window separately (see example of fluctuations in Figure 2A&B). Here we calculate the coefficient of variation (CV, the ratio of the standard deviation to the mean) for both variables for each connection to measure the fluctuation. We find that both presynaptic firing rate and synaptic efficacy fluctuate substantially over time in all areas (Figure 2C&D, CV for firing rate = 0.55±0.29 s.d., CV for efficacy = 0.69±0.46 s.d.). Although large fluctuations in firing rate are expected, large fluctuations in synaptic efficacy of this scale are somewhat unexpected during ongoing behavior. When firing rates are low, efficacy estimates are noisy, and some variation in efficacy is expected purely by chance. However, the observed fluctuations are not simply explainable by measurement error. Here we compare the observed fluctuations to surrogate data where postsynaptic spike trains are shuffled to remove any systematic fluctuations over long timescales but preserve the overall efficacy (see Methods). For each putative synapse, we shuffle 100 times to create a null distribution for CV and calculate a z-score using the mean and standard deviation of the shuffled CV. The average z-score across all putative synapses is 3.26, suggesting that the observed efficacy fluctuations are substantially higher, in most cases, than would be expected by chance. The CV for firing rate is correlated with CV for efficacy for individual synapses *r* = 0.24, *p* < 0.001. Averaging within brain regions, we also find that the CV of firing rate and the CV of efficacy are positively correlated across regions, *r* = 0.77, *p* < 0.01 (n=14 regions with >=10 putative synapses). Regions with greater fluctuations in the presynaptic firing rates tend to have higher fluctuations in the efficacies of their putative synapses.

**Figure 2.**
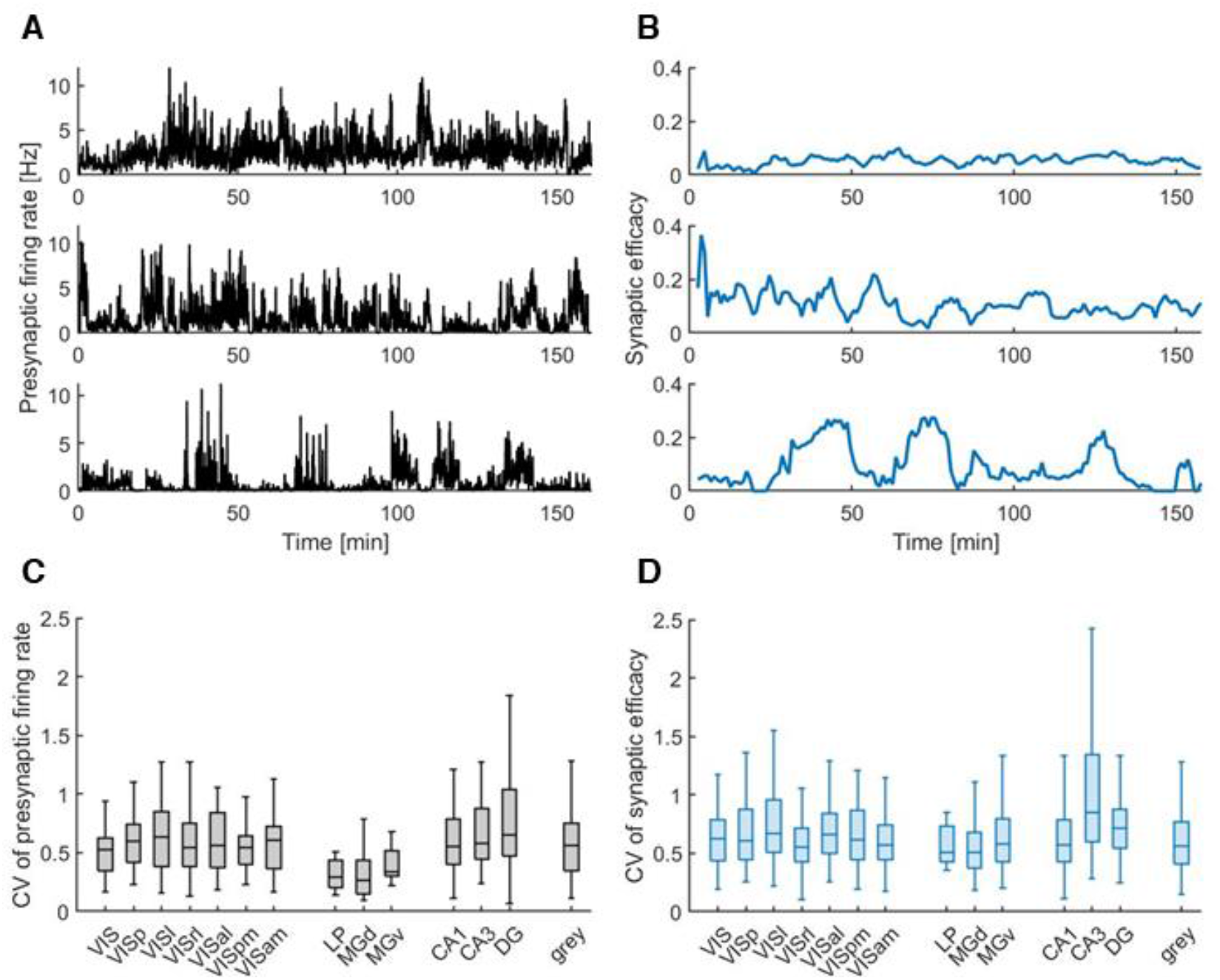
Fluctuations in presynaptic firing rate and synaptic efficacy. A & B, 3 examples for fluctuations in presynaptic firing rate (A) and synaptic efficacy (B), respectively. Top: low CV. Middle: moderate CV. Bottom: high CV. Here the firing rates in panel A are estimated using optimal time bins that are generated based on Shimazaki and Shinomoto (2010) for better visualization on finer time scales. The efficacy in panel B is estimated based on the 5-min window described in Methods. C & D, the box plots for CVs of presynaptic firing rate (C) and synaptic efficacy (D) for each area. Only areas with >=10 efferent synapses are shown here. The notations are the same as the previous figure.

### Efficacy fluctuations mirror fluctuations in the presynaptic firing rate for individual neuron pairs

The efficacies of putative synapses appear to fluctuate substantially *in vivo* across multiple brain regions. To examine these fluctuations in more detail we calculate correlations between the time-varying presynaptic firing rate and time-varying synaptic efficacy for individual connections. The average absolute Spearman correlation is 0.32±0.22 s.d. (983 out of 1383 connections are statistically significant *p*<0.05, Figure 3A & B). As with the CV of efficacy fluctuations, these correlations do not appear to be due simply to chance. By shuffling the postsynaptic spike times as described above we can also measure to what extent the Spearman correlations differ from those expected by chance. Here the average z-score of the synapses with positive correlation coefficient is 2.34, and the average z-score of the synapses with negative correlation coefficient is -1.83 (see the right panels in Figure 4 for examples). This suggests that efficacy fluctuations may be predicted based on the presynaptic firing rate, with positive or negative correlations larger than expected by chance, depending on the neuron pair. For comparison, we also examine the correlation between the fluctuations in the postsynaptic firing rate and synaptic efficacy, the average absolute Spearman correlation in this case is 0.38 ± 0.23 s.d. (1095 out of 1383 connections are significant *p*<0.05, Figure 3C & D). Positive correlations between the fluctuations in postsynaptic firing rate and synaptic efficacy are somewhat expected, since the excitability of the postsynaptic neuron may affect both (more than 75% of the synapses show a positive correlation in Figure 3C). However, the correlation between the fluctuations in the presynaptic firing rate and efficacy is less predictable and worth exploration. To further determine the influence of the presynaptic firing rate on synaptic efficacy, for each putative synapse, we run a linear regression predicting the time-varying efficacy from the log-transformed time-varying postsynaptic firing rate and the log-transformed time-varying presynaptic firing rate. Wald test shows that for 67.1% of the putative synapses, the presynaptic firing rate makes a statistically significant contribution to the prediction (928 out of 1383, *p*<0.05). For many synapses, the fluctuations in the synaptic efficacy mirror the natural fluctuations in presynaptic firing rate.

**Figure 3.**
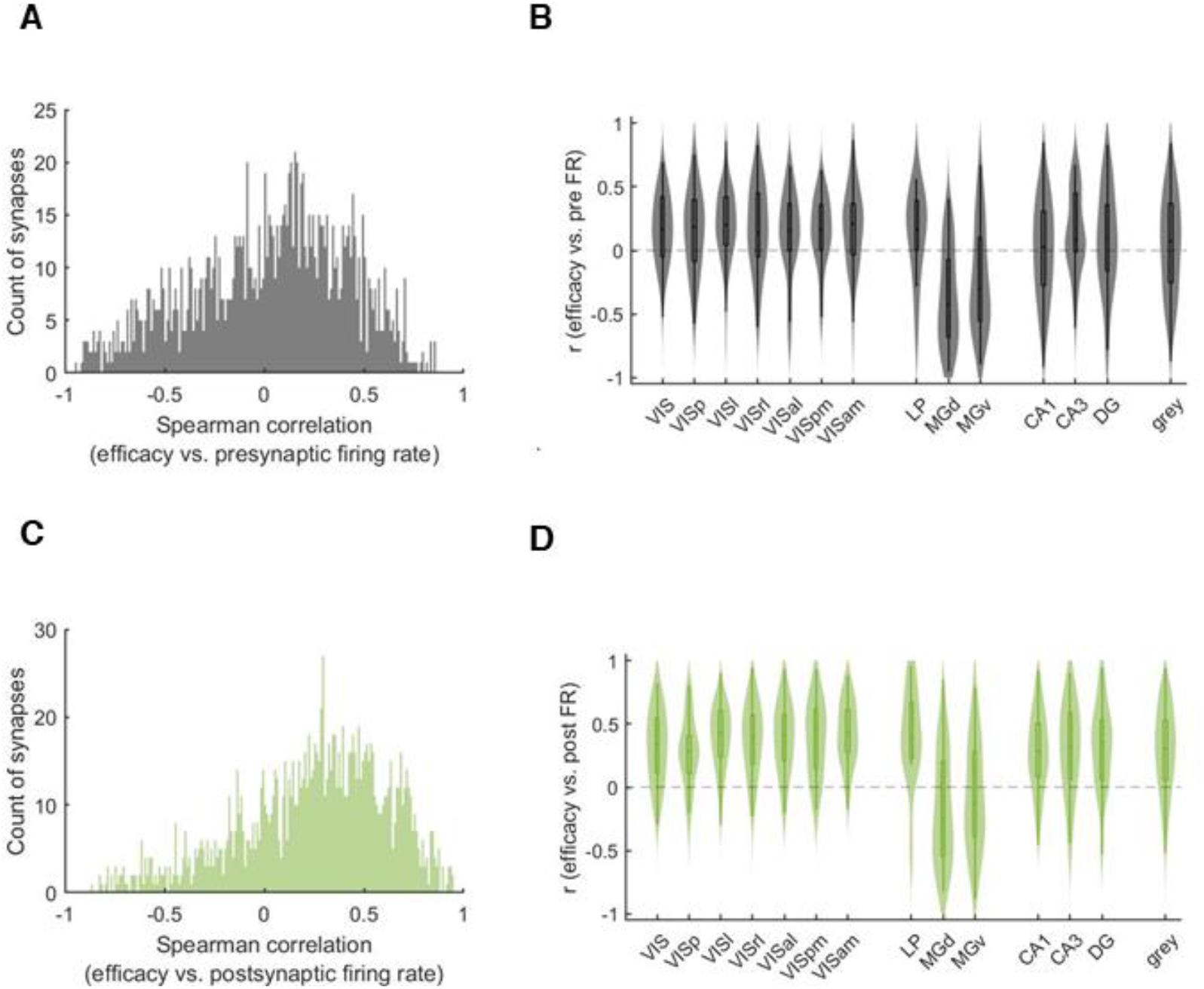
Correlation between the fluctuation in synaptic efficacy and that in the pre- and postsynaptic firing rate. A & C, the histogram of the Spearman correlation between the fluctuations in synaptic efficacy and the natural fluctuations in the presynaptic firing rate (A) or the postsynaptic firing rate (C). B & D, the violin plot of the Spearman correlation for presynaptic firing rate (B) or postsynaptic firing rate (D) for each area. Only areas with >=10 efferent synapses are shown here.

**Figure 4.**
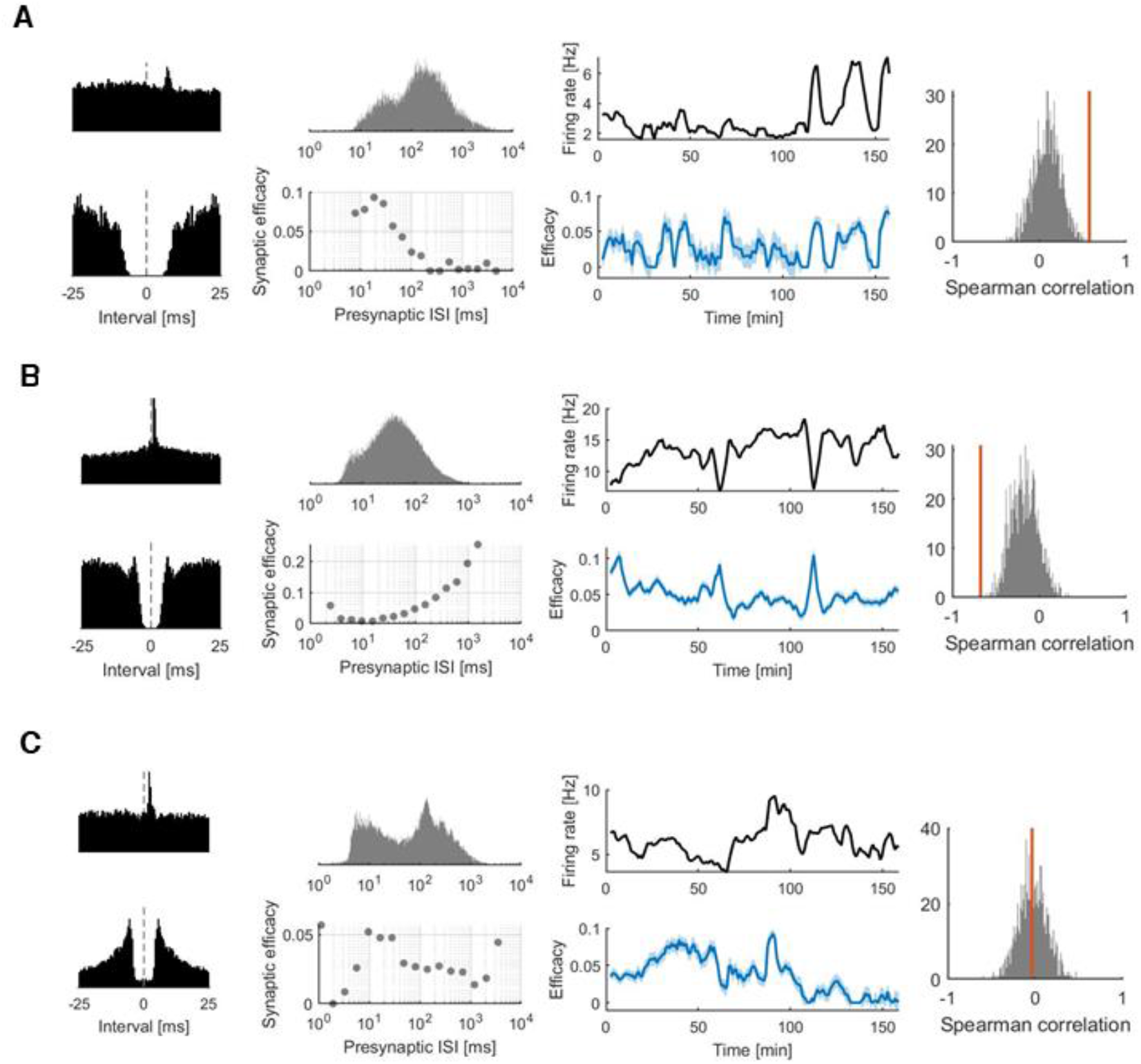
Three kinds of relationships between the fluctuations in synaptic efficacy and presynaptic firing rate. A, an example putative synapse (DG – CA3) where there is a positive correlation between the synaptic efficacy and the presynaptic firing rate. B, an example putative synapse (APN-APN) where there is a negative correlation between the synaptic efficacy and the presynaptic firing rate. C, an example putative synapse (CA1 - VIAam) where the relationship between the synaptic efficacy and the presynaptic firing rate is not clear. The first column, CCG (top) and auto-correlogram (ACG, bottom). The second column, the distribution of the presynaptic ISI (top) and the efficacy estimated using the extended GLM. The third column, the fluctuations in the presynaptic firing rate (top) and the synaptic efficacy (bottom). The shaded area represents one standard deviation. The fourth column, the distribution of the Spearman correlation from surrogate, shuffled data (grey, 1000 samples shown for visualization). The red bar shows the observed correlation from the original data. Note that in panel C, the observed correlation is close to zero. This does not mean the fluctuations in synaptic efficacy is random, it just indicates that the fluctuation in synaptic efficacy is not correlated with presynaptic firing rate.

One potential explanation for the correlation between presynaptic rate and efficacy is short-term synaptic plasticity (STP). If a synapse exhibits short-term synaptic depression higher pre-synaptic firing rates lead to a depletion of resources and lower efficacy, leading to a negative correlation. On the other hand, if a synapse exhibits short-term synaptic facilitation higher pre-synaptic rates lead to an increase in release probability and higher efficacy, leading to a positive correlation. In other cases, facilitation and depression may coexist leading to nonmonotonic relationships between presynaptic rate and efficacy. Here we find a range of relationships (Figure 4A&B) as well as cases where there are clear fluctuations but no clear relationship between presynaptic rate and efficacy (Figure 4C). This suggests that fluctuations in synaptic efficacy can potentially be predicted in detail by modeling STP.

### A model of short-term plasticity reproduces efficacy fluctuations

Since many synapses show a facilitating or depressing relationship between the presynaptic firing rate and synaptic efficacy, we examine explanatory models based on STP. Here we build a Generalized Bilinear Model (GBLM) that aims to describe the detailed (1-ms bins) postsynaptic spiking by modeling STP, slow fluctuations in postsynaptic excitability, and slow common input to both the pre- and postsynaptic neurons (see Methods). Here we estimate a “modification function” that describes how the synaptic strength changes following the occurrence of presynaptic spikes at a given ISI, and we use adaptive smoothing to track slower fluctuations in the postsynaptic rate. Altogether, this model captures the probability of postsynaptic spiking following every single presynaptic spike and allows us to track how the time-varying synaptic strength increases (facilitation) or decreases (depression) following presynaptic spikes with specific intervals.

To quantify how well the models predict the postsynaptic spiking after a presynaptic spike, we use receiver operating characteristic (ROC) curves, where the area under the curve (AUC) reflects performance (see Methods). We find that the GBLM predicts postsynaptic spikes better than the chance level (AUC = 0.68 ± 0.05 s.d.). In the 10 recordings where there is a 30-minute period of spontaneous activity, we find that the GBLM predicts postsynaptic spikes better than the chance level even when only examining this spontaneous period with we calculate the AUC using only the spikes during the spontaneous activity (AUC = 0.67 ± 0.06 s.d.). Although many putative synapses involve neurons in non-visual regions, this comparison illustrates how fluctuations in efficacy are predictable even in the absence of specific visual stimuli and not merely an artifact of fast stimulus correlations.

The GBLM provides a model of the time-varying impact of each individual presynaptic spike. However, to illustrate how slow fluctuations in efficacy might be explainable by STP more directly, we evaluate the predictions of the GBLM across 5-minute windows, as before. We find that the median coefficient of determination (*R*^2^) between the GBLM estimated efficacy and the observed efficacy is 0.22 (0.07-0.43 inter-quartile range (IQR), Figure 5A, 86.6% statistically significant, *p* < 0.05), and the results do not vary substantially across different brain regions (Figure 5B). Moreover, the modification function of presynaptic ISI from the GBLM tends to be consistent with the relationship between presynaptic firing rate and observed synaptic efficacy (Fig 5C, D, &E). As a comparison, we also build a static-synapse model without STP, where the synaptic coupling stays the same over time (see Methods). In this case, the median coefficient of determination between the static model estimated efficacy and the observed efficacy 0.17 (0.05-0.37 IQR, Figure 5A, 83.3% of them show *p* < 0.05). The static model can reproduce some fluctuations in the synaptic efficacy, indicating that some of the fluctuations can be attributed to non-STP components, such as the excitability of the postsynaptic neuron (Gal et al., 2010) or slow common inputs shared by the pre- and postsynaptic neurons. However, there is 30% of improvement in the median *R*^2^ when comparing the GBLM to the static model, which suggests that STP plays a role in the efficacy fluctuations, as well. For many connections, the GBLM can reproduce fluctuations in the synaptic efficacy with high fidelity, suggesting that STP can, at least partially, explain the relationship between fluctuations in the presynaptic firing rate and synaptic efficacy.

**Figure 5.**
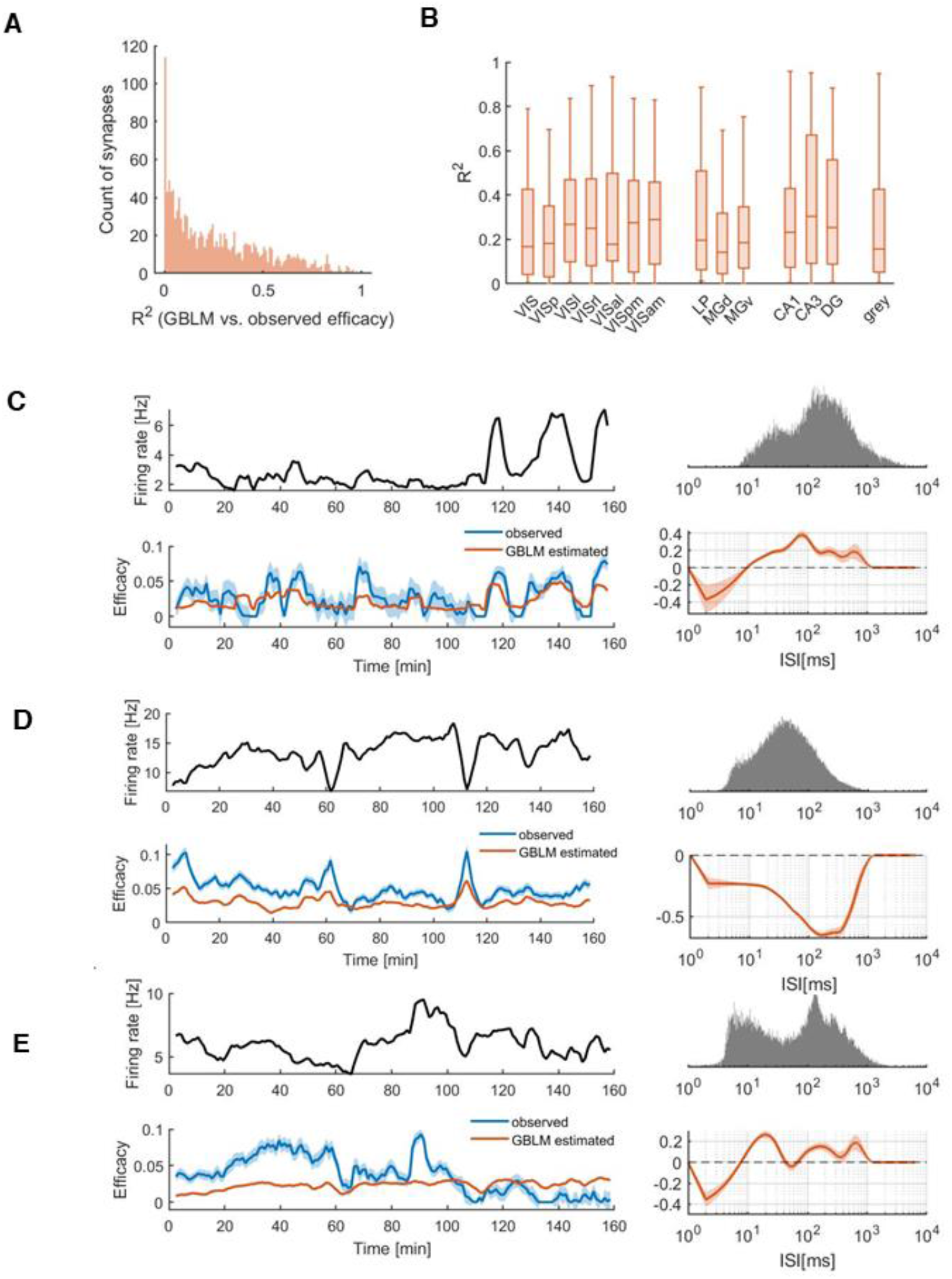
GBLM of STP reproduces the fluctuation in synaptic efficacy. **A**, the histogram of the *R*^2^ between the GBLM estimated efficacy and the observed efficacy. B, the box plot of *R*^2^ for different areas. Only areas with >=10 efferent synapses are shown here. The notations are the same as the previous figures. C & D & E, the synapses are the same ones shown in Figure 4. Left, the fluctuations in the presynaptic firing rate (top) and the synaptic efficacy (bottom). The blue line is the observed efficacy. The orange line is the GBLM estimated efficacy. Right, the distribution of the presynaptic ISI (top) and the modification function estimated from the GBLM. The shaded orange area represents 1 standard error.

To characterize the diversity of STP patterns in the putative excitatory connections examined here, we further analyze the modification functions estimated from the GBLM. The modification function gives a simple description of how the synaptic effect varies as a function of the presynaptic ISI. When the modification is positive, synaptic strength is facilitated and when it is negative it is depressed. However, the influence of the modification function, overall, depends on the ISI distribution. Even though the modification function may be negative (for ISI<10ms in Fig 5C, for example), this will only influence the synaptic strength when spikes occur with those particular ISIs. Here we align the modification function by their ISI distributions. We sample the modification function at the 5th - 95th percentiles of the ISI distribution (with 5% increment) and normalize by dividing the ISI-scaled function by the absolute value of its average. To see if there is any structure in the modification functions for all the putative synapses, we use hierarchical clustering on the ISI-scaled modification function (Euclidean distance with unweighted average linkage). We find the functions are broadly split into two clusters with some putative synapses mostly facilitating while others mostly depressing. However, the specific ISI range where facilitation or depression occurs varies widely across putative synapses (Figure 6A). Summarizing the ISI-scaled modification functions with their average, we find that the overall level of facilitation or depression appears to vary across brain regions (Figure 6B). Based on the putative synapses we detect here, the thalamic connections (originating in MGd and MGv) are dominated by depression, consistent with the observation that there are negative correlations between presynaptic rate and efficacy in these regions (Figure 3B). These results suggest that differences in the association between efficacy and presynaptic rate that we observe on ∼1minute timescales may be related to differences in STP on sub-second timescales.

**Figure 6.**
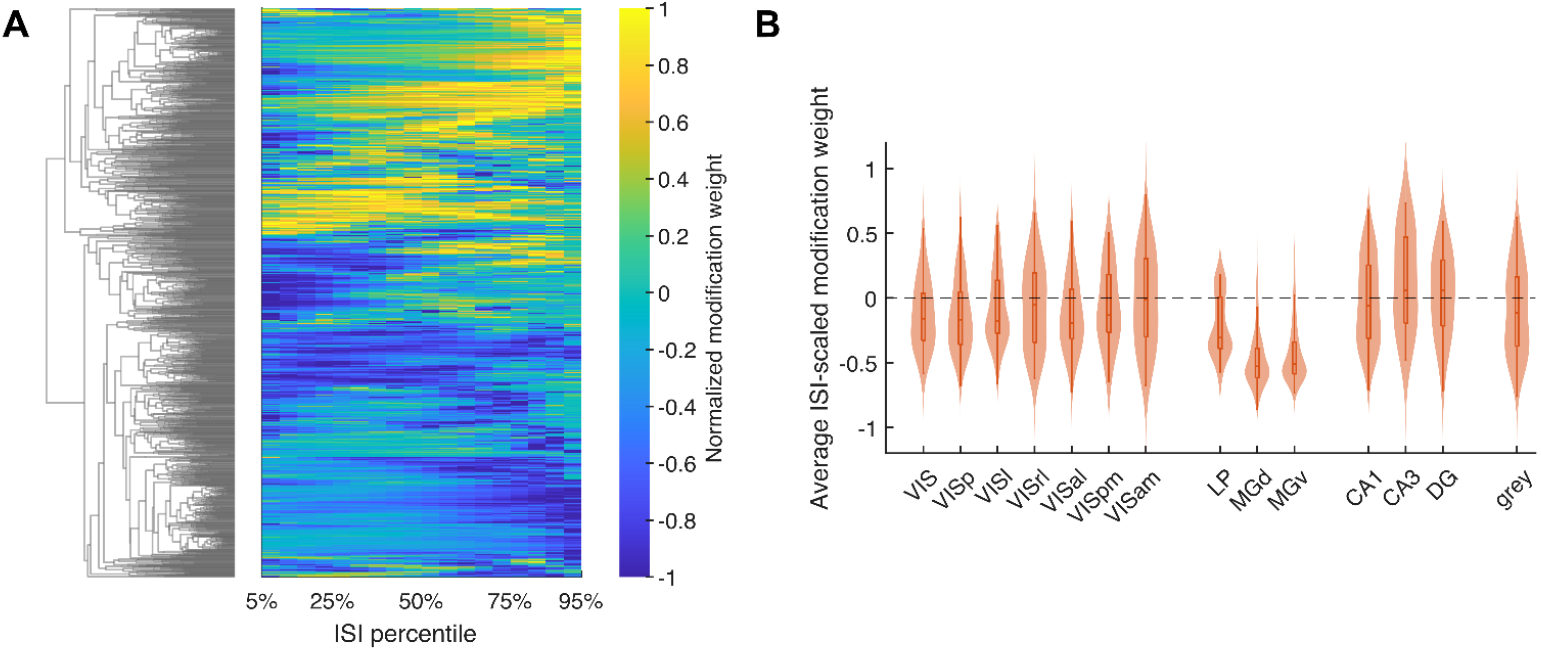
Hierarchical clustering for modification functions. A, right: the modification functions of all putative synapses sorted based on the clustering results, each row represents one modification function; left: the dendrogram of the hierarchical clustering B, the violin lot of the mean of the modification function for each brain area. Only areas with >=10 efferent synapses are shown here.

## Discussion

In this study we examine associations between fluctuations in firing rates and efficacies of putative excitatory synapses using large-scale extracellular spike recordings from awake, behaving mice. Using both descriptive and model-based analysis we identify three main findings. First, we found that, for many putative synapses, there are substantial fluctuations in the synaptic efficacy that occur on a slow timescale of minutes alongside the expected variations in pre- and postsynaptic firing rates. Second, we found that these slow fluctuations in synaptic efficacy are often correlated (either positively or negatively) with fluctuations in presynaptic rate. And third, by fitting a detailed 1ms-resolution model of postsynaptic spiking probability where synaptic effects have short-term plasticity, we show that these slow fluctuations in synaptic efficacy are consistent with the cumulative effects of STP, in many cases.

Although previous studies have shown that synaptic efficacy varies *in vivo* with learning, behavior, and stimuli (Fedulov et al., 2007; Fujisawa et al., 2008; Ghanbari et al., 2020; McKenzie et al., 2021), differences in efficacy across conditions may partially reflect differences in presynaptic firing rates and spike patterns across conditions. Firing rates depend on both external variables, such as stimuli and movement (Ben-Yishai et al., 1995; Kang et al., 2004; Vogels et al., 2005), as well as internal variables, such as arousal, attention, adaption, and excitability (Gilbert and Sigman, 2007; Gal et al., 2010; Goris et al., 2014). Firing rate fluctuations can occur spontaneously (Teich et al., 1990; Yaksi and Friedrich, 2006; Faisal et al., 2008), and are influenced by brain state and the population activity of the surrounding neurons (Tsodyks et al., 1999; Poulet and Petersen, 2008; McGinley et al., 2015). These many sources of firing rate variability occur simultaneously *in vivo* and may have complex effects on synaptic efficacy due to diversity in both cell-types and synaptic dynamics. Understanding how synaptic efficacy is affected by presynaptic firing rates, first, may allow us to better understand the influence of specific external or internal variables.

Here we find that the brain regions with larger fluctuations in the presynaptic firing rate (measured by CV) also show larger fluctuations in efficacy of putative excitatory synapses. It is important to note that this observation is just based on the putative excitatory synapses we detected. Synapse detection from spikes is biased towards strong synapses between neurons with high firing rates. Moreover, most synapses analyzed here are between neurons within the same brain area and detected on the same probe. These findings, thus, represent only a partial picture of the overall synaptic dynamics in these regions. However, the correlations we observe are consistent with our core hypothesis that firing rate variation leads directly to fluctuations in synaptic efficacy due to STP. Here we find such patterns not just across brain areas, but for many individual synapses fluctuations in synaptic efficacy mirror fluctuations in the presynaptic firing rate. Synaptic efficacy can be affected by the postsynaptic activity as well (Abbott and Nelson, 2000; Ibata et al., 2008; Sjöström et al., 2008), but here we find that the presynaptic firing rate is still predictive, even after accounting for the postsynaptic firing rate. Previous *in vitro* experiments have shown that natural spike patterns lead to changes in synaptic strength consistent with STP on short timescales (Klyachko and Stevens, 2006; Kandaswamy et al., 2010). Here, we find evidence that synaptic efficacy fluctuates with presynaptic firing rate *in vivo*, during behavior, with large fluctuations on timescales of minutes.

Using a model-based approach to predict postsynaptic spiking on 1ms-timescales (GBLM), we also examined to what extent the slow fluctuations in synaptic efficacy can be explained by STP. Here we modeled the time-varying synaptic effect using a modification function where the synaptic effect changes as a function of presynaptic ISI. We find the GBLM can predict when individual postsynaptic spikes will occur after a presynaptic spike better than chance level, as well as the accurately describe the slow fluctuations in efficacy on timescales of minutes. The modification functions estimated by the model show a range of ISI-efficacy relationships, both depressing and facilitating, and the distribution of this relationship varies in different brain areas. Previous intracellular studies have shown that short-term synaptic dynamics often depend on the pre- and postsynaptic neuron type and brain area (Thomson and Lamy, 2007; Blackman et al., 2013; Lee et al., 2019; Campagnola et al., 2022) and the presynaptic spike statistics that drive STP also vary with cell-type and brain area (Mochizuki et al., 2016). A more detailed comparison of *in vivo* results with previous *in vitro* results (Seeman et al., 2018) may help determine to what extent *in vitro* results generalize to naturalistic settings. Similarly, comparisons to previous *in vivo* studies, such as those finding depression in thalamocortical synapses, (Chung et al., 2002; Swadlow and Gusev, 2002; Bruno and Sakmann, 2006), can help determine to what extent these results generalize across species and experimental settings.

It should be noted that, by using the GBLM, we are only able to partially reproduce the fluctuations in the synaptic efficacy using the presynaptic firing activity, and many other factors could contribute to variation in synaptic efficacy. First, our model does not account for the long-term changes in the synaptic efficacy. Previous work has found that long-term plasticity can be induced in the visual cortex after repeated visual stimulation (Heynen and Bear, 2001; Frenkel et al., 2006; Cooke and Bear, 2010; Sale et al., 2011), as well as in hippocampus after behavioral training (Gruart et al., 2006; Whitlock et al., 2006; Fedulov et al., 2007). A model that accounts for the long-term plasticity may give a better prediction for the fluctuations in synaptic efficacy (Stevenson and Kording, 2011; Linderman et al., 2014; Song et al., 2018; Wei and Stevenson, 2021). Second, in our model, the STP rules themselves are assumed to be fixed for the whole recording. However, experiments have shown that the induction of long-term potentiation and long-term depression tend to shift STP itself to be more depressing or more facilitating, respectively (Markram and Tsodyks, 1996; Hardingham et al., 2007; Costa et al., 2015, 2017), which may allow for more flexibility in signal transmission (Carvalho and Buonomano, 2011). Third, although we have a term in our model to describe the slow fluctuation in network shared by the pre- and postsynaptic neurons, it does not account for the fast synaptic effect of other potential presynaptic neurons (Harris et al., 2003). A postsynaptic neuron can receive thousands of synaptic inputs, and correlated presynaptic inputs can affect the synaptic responses (Salinas and Sejnowski, 2000; Reinartz et al., 2014; Lee et al., 2016). Finally, here we model STP as a function of presynaptic ISI to directly examine the influence of presynaptic firing, but there are also many alternative models for short-term synaptic dynamics (Hennig, 2013). Our results here find that ∼20% of the variance in efficacy explained by the GBLM, but better models may be able to improve on this result and more accurately predict fluctuations in synaptic efficacy *in vivo*.

Using extracellular spike recordings from behaving animals, here we find that the substantial fluctuations in the efficacy of putative excitatory synapses can be partially predicted by modeling short-term synaptic plasticity. Large-scale spike recordings have the advantage of allowing us to study putative connections between many brain areas for long periods of time (>2hrs). Data from a wide range of synapses across the entire brain, may drive richer theoretical and normative models of synaptic transmission (Pfister et al., 2010; Hennig, 2013). However, studying synapses from spikes has the disadvantage that it requires sufficient numbers of presynaptic spikes to accurately estimate efficacy. Advances in stimulation and recording techniques may make it possible to study fluctuations on faster timescales (Kwan and Dan, 2012; English et al., 2017; Walker et al., 2021; Hage et al., 2022), and systematic fluctuations in efficacy for inhibitory synapses may also occur (Reyes et al., 1998; Beierlein et al., 2003; Ma et al., 2012). Although disentangling the complex relationships between synaptic efficacy and external or internal variables *in vivo* is an ongoing challenge in systems neuroscience, our results here highlight the potentially important role of STP in driving slow fluctuations in synaptic efficacy. In putative excitatory synapses from a wide range of brain areas, we find that short-term synaptic dynamics may lead to large, predictable fluctuations on longer timescales.

## Code Accessibility

The MATLAB code for data analysis is available at GitHub: https://github.com/NaixinRen/predictable-fluctuations-in-synaptic-efficacy.

## Acknowledgements

This material is based upon work supported by the National Science Foundation under Grant No. 1651396. We thank Harvey Swadlow and Monty Escabi for helpful discussions and would also like to thank the Allen Institute for Brain Science for sharing the Visual Coding – Neuropixels dataset and for supporting open science.

## Notes

### Competing Interest Statement

The authors have declared no competing interest.

## References

Abbott LF, Nelson SB (2000) Synaptic plasticity: Taming the beast. Nat Neurosci 3:1178–1183 Available at: https://www.nature.com/articles/nn1100_1178 [Accessed November 21, 2021].

Abbott LF, Varela JA, Sen K, Nelson SB (1997) Synaptic depression and cortical gain control. Science (80-) 275:220–224 Available at: http://science.sciencemag.org/ [Accessed November 12, 2020].

Amarasingham A, Harrison MT, Hatsopoulos NG, Geman S (2012) Conditional modeling and the jitter method of spike resampling. J Neurophysiol 107:517–531 Available at: https://journals.physiology.org/doi/abs/10.1152/jn.00633.2011 [Accessed February 3, 2022].

Barthó P, Hirase H, Monconduit L, Zugaro M, Harris KD, Buzsáki G (2004) Characterization of neocortical principal cells and interneurons by network interactions and extracellular features. J Neurophysiol 92:600–608.

Beierlein M, Gibson JR, Connors BW (2003) Two Dynamically Distinct Inhibitory Networks in Layer 4 of the Neocortex. J Neurophysiol 90:2987–3000 Available at: https://journals.physiology.org/doi/full/10.1152/jn.00283.2003 [Accessed April 13, 2022].

Ben-Yishai R, Lev Bar-Or R, Sompolinsky H (1995) Theory of orientation tuning in visual cortex. Proc Natl Acad Sci U S A 92:3844–3848 Available at: https://www.pnas.org [Accessed March 6, 2022].

Blackman A V., Abrahamsson T, Costa RP, Lalanne T, Sjöström PJ (2013) Target-cell-specific short-term plasticity in local circuits. Front Synaptic Neurosci 5:1–13.

Bruno RM, Sakmann B (2006) Cortex is driven by weak but synchronously active thalamocortical synapses. Science (80-) 312:1622–1627 Available at: https://www.science.org [Accessed March 5, 2022].

Buzsáki G, Mizuseki K (2014) The log-dynamic brain: How skewed distributions affect network operations. Nat Rev Neurosci 15:264–278 Available at: https://www.nature.com/articles/nrn3687 [Accessed February 17, 2022].

Campagnola L et al. (2022) Local connectivity and synaptic dynamics in mouse and human neocortex. Science (80-) 375 Available at: https://www.science.org/doi/full/10.1126/science.abj5861 [Accessed April 14, 2022].

Carvalho TP, Buonomano D V. (2011) A novel learning rule for long-term plasticity of short-term synaptic plasticity enhances temporal processing. Front Integr Neurosci 5:20.

Chung S, Li X, Nelson SB (2002) Short-term depression at thalamocortical synapses contributes to rapid adaptation of cortical sensory responses in vivo. Neuron 34:437–446 Available at: https://www.sciencedirect.com/science/article/pii/S0896627302006591 [Accessed December 14, 2020].

Churchland MM, Cunningham JP, Kaufman MT, Foster JD, Nuyujukian P, Ryu SI, Shenoy K V., Shenoy K V. (2012) Neural population dynamics during reaching. Nature 487:51–56 Available at: https://www.nature.com/articles/nature11129 [Accessed October 16, 2020].

Connors BW, Gutnick MJ (1990) Intrinsic firing patterns of diverse neocortical neurons. Trends Neurosci 13:99–104.

Cooke SF, Bear MF (2010) Visual experience induces long-term potentiation in the primary visual cortex. J Neurosci 30:16304–16313 Available at: https://www.jneurosci.org/content/30/48/16304 [Accessed March 5, 2022].

Costa RP, Froemke RC, Sjöström PJ, van Rossum MCW (2015) Unified pre- and postsynaptic long-term plasticity enables reliable and flexible learning. Elife 4:1–16 Available at: https://elifesciences.org/articles/9457 [Accessed February 17, 2020].

Costa RP, Jesper Sjöström P, van Rossum MCW (2013) Probabilistic inference of short-term synaptic plasticity in neocortical microcircuits. Front Comput Neurosci 7:75 Available at: http://journal.frontiersin.org/article/10.3389/fncom.2013.00075/abstract [Accessed February 17, 2020].

Costa RP, Mizusaki BEP, Sjöström PJ, van Rossum MCW (2017) Functional consequences of pre- and postsynaptic expression of synaptic plasticity. Philos Trans R Soc B Biol Sci 372:20160153 Available at: http://biorxiv.org/content/early/2016/10/29/075317.abstract [Accessed February 17, 2020].

Deng PY, Klyachko VA (2011) The diverse functions of short-term plasticity components in synaptic computations. Commun Integr Biol 4:543–548 Available at: https://www.tandfonline.com/doi/abs/10.4161/cib.15870 [Accessed March 21, 2022].

English DF, Mckenzie S, Evans T, Kim K, Yoon E, Buzsáki G (2017) Pyramidal cell-interneuron circuit architecture and dynamics in hippocampal networks. :505–520.

Faisal AA, Selen LPJ, Wolpert DM (2008) Noise in the nervous system. Nat Rev Neurosci 9:292–303 Available at: https://www.nature.com/articles/nrn2258 [Accessed March 2, 2022].

Fedulov V, Rex CS, Simmons DA, Palmer L, Gall CM, Lynch G (2007) Evidence that long-term potentiation occurs within individual hippocampal synapses during learning. J Neurosci 27:8031–8039 Available at: https://www.jneurosci.org/content/27/30/8031 [Accessed March 5, 2022].

Fetz E, Toyama K, Smith W (1991) Synaptic Interactions between Cortical Neurons. In: Normal and Altered States of Function. Cerebral cortex (Peters A, Jones EG, eds), pp 1–47. Boston, MA: Springer. Available at: https://link.springer.com/chapter/10.1007/978-1-4615-6622-9_1 [Accessed October 23, 2019].

Fortune ES, Rose GJ (2001) Short-term synaptic plasticity as a temporal filter. Trends Neurosci 24:381–385 Available at: https://www.sciencedirect.com/science/article/pii/S016622360001835X?casa_token=suI0FzRCgTwAAAAA:YXny16xdLvhicL8MFB17U0UzH5nJpB5KHs_0V62jd0UiPtnQ0E00FGLHTdAmaXhyyOS9tKV_Xw [Accessed December 15, 2020].

Frenkel MY, Sawtell NB, Diogo ACM, Yoon B, Neve RL, Bear MF (2006) Instructive Effect of Visual Experience in Mouse Visual Cortex. Neuron 51:339–349 Available at: https://www.sciencedirect.com/science/article/pii/S089662730600506X [Accessed March 5, 2022].

Fujisawa S, Amarasingham A, Harrison MT, Buzsáki G (2008) Behavior-dependent short-term assembly dynamics in the medial prefrontal cortex. Nat Neurosci.

Gal A, Eytan D, Wallach A, Sandler M, Schiller J, Marom S (2010) Dynamics of excitability over extended timescales in cultured cortical neurons. J Neurosci 30:16332–16342 Available at: https://www.jneurosci.org/content/30/48/16332 [Accessed April 13, 2022].

Ghanbari A, Malyshev A, Volgushev M, Stevenson IH (2017) Estimating short-term synaptic plasticity from pre- and postsynaptic spiking. PLOS Comput Biol 13:e1005738 Available at: https://doi.org/10.1371/journal.pcbi.1005738.

Ghanbari A, Ren N, Keine C, Stoelzel C, Englitz B, Swadlow HA, Stevenson IH (2020) Modeling the short-term dynamics of in vivo excitatory spike transmission. J Neurosci 40:4185–4202 Available at: https://doi.org/10.1523/JNEUROSCI.1482-19.2020 [Accessed September 9, 2020].

Gilbert CD, Sigman M (2007) Brain States: Top-Down Influences in Sensory Processing. Neuron 54:677–696.

Goris RLT, Movshon JA, Simoncelli EP (2014) Partitioning neuronal variability. Nat Neurosci 17:858–865 Available at: https://www.nature.com/articles/nn.3711 [Accessed March 1, 2022].

Gruart A, Muñoz MD, Delgado-García JM (2006) Involvement of the CA3-CA1 synapse in the acquisition of associative learning in behaving mice. J Neurosci 26:1077–1087 Available at: https://www.jneurosci.org/content/26/4/1077.short [Accessed March 5, 2022].

Hage TA, Bosma-Moody A, Baker CA, Kratz M, Campagnola L, Jarsky T, Zeng H, Murphy GJ (2022) Synaptic connectivity to L2/3 of primary visual cortex measured by two-photon optogenetic stimulation. Elife 11.

Hardingham NR, Hardingham GE, Fox KD, Jack JJB (2007) Presynaptic efficacy directs normalization of synaptic strength in layer 2/3 rat neocortex after paired activity. J Neurophysiol 97:2965–2975 Available at: https://journals.physiology.org/doi/abs/10.1152/jn.01352.2006 [Accessed March 5, 2022].

Harris KD, Csicsvari J, Hirase H, Dragoi G, Buzsáki G (2003) Organization of cell assemblies in the hippocampus. Nature 424:552–556.

Hennig MH (2013) Theoretical models of synaptic short term plasticity. Front Comput Neurosci 7:45.

Heynen AJ, Bear MF (2001) Long-term potentiation of thalamocortical transmission in the adult visual cortex in vivo. J Neurosci 21:9801–9813 Available at: https://www.jneurosci.org/content/21/24/9801 [Accessed March 5, 2022].

Ibata K, Sun Q, Turrigiano GG (2008) Rapid Synaptic Scaling Induced by Changes in Postsynaptic Firing. Neuron 57:819–826.

Kandaswamy U, Deng PY, Stevens CF, Klyachko VA (2010) The role of presynaptic dynamics in processing of natural spike trains in hippocampal synapses. J Neurosci 30:15904–15914 Available at: www.jneurosci.org [Accessed August 21, 2020].

Kang K, Shapley RM, Sompolinsky H (2004) Information Tuning of Populations of Neurons in Primary Visual Cortex. J Neurosci 24:3726–3735 Available at: https://www.jneurosci.org/content/24/15/3726 [Accessed March 6, 2022].

Klyachko VA, Stevens CF (2006) Excitatory and feed-forward inhibitory hippocampal synapses work synergistically as an adaptive filter of natural spike trains Segev I, ed. PLoS Biol 4:1187–1200 Available at: https://dx.plos.org/10.1371/journal.pbio.0040207 [Accessed August 21, 2020].

Kobayashi R, Kurita S, Kurth A, Kitano K, Mizuseki K, Diesmann M, Richmond BJ, Shinomoto S (2019) Reconstructing neuronal circuitry from parallel spike trains. Nat Commun 10.

Kwan AC, Dan Y (2012) Dissection of cortical microcircuits by single-neuron stimulation in vivo. Curr Biol 22:1459–1467.

Lee JH, Campagnola L, Seeman SC, Jarsky T, Mihalas S (2019) Functional synapse types via characterization of short-term synaptic plasticity. bioRxiv: 10.1101/648725 Available at: https://www.biorxiv.org/content/10.1101/648725v1.

Lee KFH, Soares C, Thivierge JP, Béïque JC (2016) Correlated Synaptic Inputs Drive Dendritic Calcium Amplification and Cooperative Plasticity during Clustered Synapse Development. Neuron 89:784–799.

Levick WR, Cleland BG, Dubin MW (1972) Lateral geniculate neurons of cat: retinal inputs and physiology. Invest Ophthalmol 11:302–311 Available at: https://www.academia.edu/download/42823304/Lateral_geniculate_neurons_of_the_cat_re20160218-31272-1wqevzp.pdf [Accessed March 14, 2022].

Linderman SW, Stock CH, Adams RP (2014) A framework for studying synaptic plasticity with neural spike train data. In: Advances in Neural Information Processing Systems, pp 2330–2338.

Ma Y, Hu H, Agmon A (2012) Short-term plasticity of unitary inhibitory-to-inhibitory synapses depends on the presynaptic interneuron subtype. J Neurosci 32:983–988 Available at: https://www.jneurosci.org/content/32/3/983 [Accessed April 13, 2022].

Markram H, Tsodyks M (1996) Redistribution of synaptic efficacy between neocortical pyramidal neurons. Nature 382:807–810 Available at: https://www.nature.com/articles/382807a0 [Accessed February 17, 2020].

McGinley MJ, Vinck M, Reimer J, Batista-Brito R, Zagha E, Cadwell CR, Tolias AS, Cardin JA, McCormick DA (2015) Waking State: Rapid Variations Modulate Neural and Behavioral Responses. Neuron 87:1143–1161.

McKenzie S, Huszár R, English DF, Kim K, Christensen F, Yoon E, Buzsáki G (2021) Preexisting hippocampal network dynamics constrain optogenetically induced place fields. Neuron 109:1040-1054.e7.

Mochizuki Y et al. (2016) Similarity in neuronal firing regimes across mammalian species. J Neurosci 36:5736–5747 Available at: https://www.jneurosci.org/content/36/21/5736 [Accessed February 3, 2022].

Pala A, Petersen CCH (2015) InVivo Measurement of Cell-Type-Specific Synaptic Connectivity and Synaptic Transmission in Layer 2/3 Mouse Barrel Cortex. Neuron 85:68–75.

Pala A, Petersen CCH (2018) State-dependent cell-type-specific membrane potential dynamics and unitary synaptic inputs in awake mice. Elife 7.

Perkel DH, Gerstein GL, Moore GP (1967) Neuronal Spike Trains and Stochastic Point Processes: II. Simultaneous Spike Trains. Biophys J 7:419–440 Available at: https://www.sciencedirect.com/science/article/pii/S0006349567865974 [Accessed October 3, 2019].

Pfister JP, Dayan P, Lengyel M (2010) Synapses with short-term plasticity are optimal estimators of presynaptic membrane potentials. Nat Neurosci 13:1271–1275 Available at: https://www.nature.com/articles/nn.2640 [Accessed March 6, 2022].

Pillow JW, Shlens J, Paninski L, Sher A, Litke AM, Chichilnisky EJ, Simoncelli EP (2008) Spatio-temporal correlations and visual signalling in a complete neuronal population. Nature 454:995–999 Available at: https://www.nature.com/articles/nature07140 [Accessed October 15, 2020].

Poulet JFA, Petersen CCH (2008) Internal brain state regulates membrane potential synchrony in barrel cortex of behaving mice. Nature 454:881–885 Available at: https://www.nature.com/articles/nature07150 [Accessed March 2, 2022].

Rebesco JM, Stevenson IH, Körding KP, Solla SA, Miller LE (2010) Rewiring neural interactions by micro-stimulation. Front Syst Neurosci 4:39.

Reinartz S, Biro I, Gal A, Giugliano M, Marom S (2014) Synaptic dynamics contribute to long-term single neuron response fluctuations. Front Neural Circuits 8:71.

Ren N, Ito S, Hafizi H, Beggs JM, Stevenson IH (2020) Model-based detection of putative synaptic connections from spike recordings with latency and type constraints. J Neurophysiol Available at: https://doi.org/10.1152/jn.00066.2020.

Reyes A, Lujan R, Rozov A, Burnashev N, Somogyi P, Sakmann B (1998) Target-cell-specific facilitation and depression in neocortical circuits. Nat Neurosci 1:279–284 Available at: https://www.nature.com/articles/nn0898_279 [Accessed April 13, 2022].

Sale A, De Pasquale R, Bonaccorsi J, Pietra G, Olivieri D, Berardi N, Maffei L (2011) Visual perceptual learning induces long-term potentiation in the visual cortex. Neuroscience 172:219–225.

Salinas E, Sejnowski TJ (2000) Impact of correlated synaptic input on output firing rate and variability in simple neuronal models. J Neurosci 20:6193–6209 Available at: https://www.jneurosci.org/content/20/16/6193 [Accessed March 5, 2022].

Sedigh-Sarvestani M, Vigeland L, Fernandez-Lamo I, Taylor MM, Palmer LA, Contreras D (2017) Intracellular, in vivo, dynamics of thalamocortical synapses in visual cortex. J Neurosci 37:5250–5262 Available at: https://www.jneurosci.org/content/37/21/5250 [Accessed April 13, 2022].

Seeman SC et al. (2018) Sparse recurrent excitatory connectivity in the microcircuit of the adult mouse and human cortex. Elife 7.

Shenoy K V., Sahani M, Churchland MM (2013) Cortical control of arm movements: A dynamical systems perspective. Annu Rev Neurosci 36:337–359 Available at: http://www.annualreviews.org/doi/10.1146/annurev-neuro-062111-150509 [Accessed November 20, 2020].

Shimazaki H, Shinomoto S (2010) Kernel bandwidth optimization in spike rate estimation. J Comput Neurosci 29:171–182 Available at: https://link.springer.com/article/10.1007/s10827-009-0180-4 [Accessed November 22, 2021].

Siegle JH et al. (2021) Survey of spiking in the mouse visual system reveals functional hierarchy. Nature 592:86–92 Available at: https://www.nature.com/articles/s41586-020-03171-x [Accessed November 21, 2021].

Sjöström PJ, Rancz EA, Roth A, Häusser M (2008) Dendritic excitability and synaptic plasticity. Physiol Rev 88:769–840 Available at: https://journals.physiology.org/doi/abs/10.1152/physrev.00016.2007 [Accessed March 5, 2022].

Song D, Robinson BS, Berger TW (2018) Identification of short-term and long-term functional synaptic plasticity from spiking activities. In: Adaptive Learning Methods for Nonlinear System Modeling, pp 289–312. Elsevier. Available at: https://www.sciencedirect.com/science/article/pii/B9780128129760000178 [Accessed March 5, 2022].

Stevenson IH, Kording KP (2011) Inferring spike-timing-dependent plasticity from spike train data. In: Advances in Neural Information Processing Systems 24: 25th Annual Conference on Neural Information Processing Systems 2011, NIPS 2011 (Shawe-Taylor J, Zemel RS, Bartlett P, Pereira FCN, Weinberger KQ, eds), pp 1–9 Available at: https://proceedings.neurips.cc/paper/2011/hash/602d1305678a8d5fdb372271e980da6a-Abstract.html [Accessed March 5, 2022].

Stoelzel CR, Bereshpolova Y, Gusev AG, Swadlow HA (2008) The impact of an LGNd impulse on the awake visual cortex: Synaptic dynamics and the sustained/transient distinction. J Neurosci 28:5018–5028.

Stoelzel CR, Bereshpolova Y, Swadlow HA (2009) Stability of thalamocortical synaptic transmission across awake brain states. J Neurosci 29:6851–6859.

Stoelzel CR, Huff JM, Bereshpolova Y, Zhuang J, Hei X, Alonso JM, Swadlow HA (2015) Hour-long adaptation in the awake early visual system. J Neurophysiol 114:1172–1182 Available at: https://journals.physiology.org/doi/abs/10.1152/jn.00116.2015 [Accessed January 11, 2022].

Swadlow HA, Gusev AG (2001) The impact of “bursting” thalamic impulses at a neocortical synapse. Nat Neurosci 4:402–408 Available at: https://www.nature.com/articles/nn0401_402 [Accessed March 14, 2022].

Swadlow HA, Gusev AG (2002) Receptive-field construction in cortical inhibitory interneurons. Nat Neurosci 5:403–404.

Teich MC, Johnson DH, Kumar AR, Turcott RG (1990) Rate fluctuations and fractional power-law noise recorded from cells in the lower auditory pathway of the cat. Hear Res 46:41–52.

Thomson AM, Lamy C (2007) Functional Maps of Neocortical Local Circuitry. Front Neurosci 1:19–42.

Trainito C, von Nicolai C, Miller EK, Siegel M (2019) Extracellular Spike Waveform Dissociates Four Functionally Distinct Cell Classes in Primate Cortex. Curr Biol 29:2973-2982.e5.

Truccolo W, Eden UT, Fellows MR, Donoghue JP, Brown EN (2005) A point process framework for relating neural spiking activity to spiking history, neural ensemble, and extrinsic covariate effects. J Neurophysiol 93:1074–1089.

Tsodyks M, Kenet T, Grinvald A, Arieli A (1999) Linking spontaneous activity of single cortical neurons and the underlying functional architecture. Science (80-) 286:1943–1946 Available at: https://www.science.org/doi/abs/10.1126/science.286.5446.1943 [Accessed March 2, 2022].

Tsodyks M V., Markram H (1997) The neural code between neocortical pyramidal neurons depends on neurotransmitter release probability. Proc Natl Acad Sci U S A 94:719–723.

Usrey WM, Alonso JM, Reid RC (2000) Synaptic interactions between thalamic inputs to simple cells in cat visual cortex. J Neurosci 20:5461–5467 Available at: https://www.jneurosci.org/content/20/14/5461 [Accessed March 14, 2022].

Vogels TP, Rajan K, Abbott LF (2005) Neural network dynamics. Annu Rev Neurosci 28:357–376 Available at: https://www.annualreviews.org/doi/abs/10.1146/annurev.neuro.28.061604.135637 [Accessed March 6, 2022].

Walker AS, Raliski BK, Karbasi K, Zhang P, Sanders K, Miller EW (2021) Optical Spike Detection and Connectivity Analysis With a Far-Red Voltage-Sensitive Fluorophore Reveals Changes to Network Connectivity in Development and Disease. Front Neurosci 15:352.

Wei G, Stevenson IH (2021) Tracking fast and slow changes in synaptic weights from simultaneously observed pre- and postsynaptic spiking. Neural Comput 33:2682–2709 Available at: https://arxiv.org/abs/2102.01803v2 [Accessed November 16, 2021].

Whitlock JR, Heynen AJ, Shuler MG, Bear MF (2006) Learning induces long-term potentiation in the hippocampus. Science (80-) 313:1093–1097 Available at: https://science.sciencemag.org/content/313/5790/1093.abstract [Accessed November 21, 2021].

Yaksi E, Friedrich RW (2006) Reconstruction of firing rate changes across neuronal populations by temporally deconvolved Ca2+ imaging. Nat Methods 3:377–383 Available at: https://www.nature.com/articles/nmeth874 [Accessed March 2, 2022].

Zucker RS, Regehr WG (2002) Short-term synaptic plasticity. Annu Rev Physiol 64:355–405 Available at: https://www.annualreviews.org/doi/abs/10.1146/annurev.physiol.64.092501.114547 [Accessed November 21, 2021].

